# Paying Attention to Other Animal Detections Improves Camera Trap Classification Models

**DOI:** 10.1101/2025.07.15.664849

**Authors:** Gaspard Dussert, Stéphane Dray, Simon Chamaillé-Jammes, Vincent Miele

## Abstract

1. In ecological studies, automated species classification models are increasingly used to process large volumes of camera trap images. Most current classification models rely on a two-step pipeline: a detector first locates and crops animals, followed by a classifier that predicts species independently for each crop. While effective, these models still struggle under challenging conditions and ignore temporal context or information from nearby animals available in sequences of camera trap images, whereas human annotators often use it in difficult cases.
2. We propose to leverage self-attention, a core mechanism of Large Language Models and Vision Transformers, to enable the model to learn relationships between crops occurring in similar contexts. Our self-attention module operates directly on the set of crop embeddings, producing new representations enriched with information from other crops, improving species classification. The module fits into the two-step pipeline without requiring structural change and adds only minimal computational overhead. To address the lack of annotated multi-species sequences, we develop a training strategy that synthetically generates multi-species sequences from monospecific ones.
3. Compared to an independent classifier baseline, our multi-crop model achieves higher accuracy on mono-specific sequences, both real and synthetic. In multispecies settings, evaluated using synthetic test sets, we also observe a substantial improvement in accuracy. Using real but weakly annotated multi-species sequences, we reformulate the task as multi-label set classification, and conduct a visual analysis, to highlight the benefits of our approach.
4. By leveraging the information brought by all detections of animals in the image and others of the same sequence, our approach reduces species misclassifications and enables more accurate estimates for downstream ecological analyses focusing on, for instance, species richness, occupancy, and species interaction.

## 1 Introduction

Camera traps have become essential tools in wildlife monitoring, enabling large-scale and non-invasive data collection across a wide range of ecosystems. Their growing use brings the challenge of identifying the species present in thousands to millions of images or videos. As manual identification is time-consuming and tedious, developing efficient and accurate automated analysis pipelines is much needed (Tuia et al. 2022; Pollock et al. 2025). Most modern camera trap analysis pipelines now adopt a two-step procedure with two deep-learning models: an object detector (e.g. MegaDetector v5 or v6, Beery et al. 2019a; Hernandez et al. 2024) first detects animals within images, and then a species classifier is applied to each animal cropped image (hereafter referred to as crop; Norman et al. 2023; Gadot et al. 2024). This approach is now well-established and implemented in several widely adopted popular tools, software, and web platforms (Lunteren 2023; Hernandez et al. 2024; Rigoudy et al. 2023; Osner 2024; Ahumada et al. 2020). It is also particularly well suited to multi-species sequences, as each crop is processed independently by the classifier, allowing multiple species to be predicted from the same image.

Researchers now regularly train species-classification deep-learning models that often exceed 90% accuracy (Norouzzadeh et al. 2018; Whytock et al. 2021; Willi et al. 2019) but higher results seem to be complex to achieve. They still struggle with challenging conditions, such as motion blur, occlusion, low image quality, or unusual animal poses. These errors can propagate and negatively affect downstream ecological tasks, such as estimating species richness, occupancy, or interspecies interactions (Sollmann 2018; Gimenez et al. 2022; Nicvert et al. 2024).

Predicting each animal crop independently increases the likelihood of errors, especially for animals further from the camera trap, whose crops tend to be of lower quality. For this reason, human annotators frequently rely on the information accumulated elsewhere to make more accurate species identifications. First, since camera traps often capture short sequences of consecutive images (aka events, bursts, or episodes), human annotators frequently use the temporal context to make more accurate species identifications. This is particularly true when key identifying features are missing in some images of the sequence but clearly visible in others. Second, since many social species are photographed in groups, human annotators use this information to confirm the identification of an individual based on the other individuals in a group. However, this additional information is overlooked by existing classification models.

A straightforward way to incorporate temporal context or information from surrounding animals is to assign each individual crop the best prediction (or the average prediction) across the image or the sequence. Such approaches have shown good performance, but they assume that only a single species is present (Dussert et al. 2024; Shashidhara et al. 2020; Willi et al. 2019). While multi-species sequences are rare in some habitats, failing to detect them can still be problematic, as these co-occurrences often correspond to ecologically significant interspecies interactions. In other habitats, multi-species sequences are much more frequent, and methods assuming a single species would lead to many classification errors. While several prior works have explored the use of temporal or contextual information in camera trap data (Beery et al. 2019b; Yang et al. 2019), these methods fall outside the established two-step pipeline, limiting their adoption. Moreover, they primarily focus on improving detection rather than species classification. While video models, such as TimeSformer (Bertasius et al. 2021), can leverage temporal cues between frames, they are ill-suited for our task: unlikes videos, the sets of crops we aim to process are neither temporally coherent nor spatially consistent.

Therefore, to replicate the human ability to connect information across different parts of an image or sequence, automated systems must include a mechanism capable of learning relationships between objects (here, the crops). In recent years, the self-attention mechanism has gained popularity through its central role in large language models (Vaswani et al. 2017). Self-attention is a trainable mechanism that learns to capture dependencies and contextual links between elements in a sequence. In language models, it operates on words or tokens, and in state-of-the-art vision models like Vision Transformers (ViT), it is applied to image patches (Dosovitskiy et al. 2021). We propose to leverage this principle to model relationships between crops in camera trap sequences.

In this work, our goal is to enhance species classification within the two-step framework without requiring fundamental changes to its structure or compromising performance on multi-species sequences. We aim to build on existing component, such as MegaDetector (Beery et al. 2019a; Hernandez et al. 2024) and pretrained species classifiers, avoiding the need for labour-intensive annotations of animal bounding boxes, and allowing the use of high-performing classification models such as EfficientNetV2 (Tan et al. 2021) or DINOv2 (Oquab et al. 2023). Our key idea is to propose adding a lightweight module based on self-attention. In our case, we apply self-attention to the set of all embeddings of animal crops detected within the same camera trap sequence, allowing the model to capture relationships between them. Here, an embedding refers to a high-dimensional feature vector produced by a neural network, capturing semantic information about each crop. Self-attention allows each crop embedding to be refined by incorporating contextual information from the full set of crops in the image or sequence. Importantly, this module adds minimal computational overhead at inference time and can be seamlessly integrated into existing pipelines. We design a training recipe to effectively train such a self-attention module for our specific use case. Due to the absence of multi-species sequences annotated at the crop level, we introduce a method to generate synthetic sets of crops by combining mono-specific ones. This approach enables the model to be trained with more diverse data and enables the model to learn how to handle multi-specific events.

## 2 Material and Methods

### 2.1 Data

To evaluate our method, we use two datasets publicly available in the LILA repository (https://lila.science/).

#### Snapshot Serengeti

This large-scale camera trap dataset collected in *Serengeti* National Park between 2010 and 2018 (Swanson et al. 2015). It comprises 7.18 million images from 2.66 million sequences (i.e., sets of consecutive images) captured at 225 camera trap locations. Annotations are provided at the sequence level, meaning that labels apply to the entire sequence rather than to individual images or animals. The dataset includes 61 categories (species and higher taxonomic groups). 4.37% of the non-empty sequences are annotated with multiple labels (e.g a whole sequence is labelled zebra and wildebeest). Throughout this paper, we refer to this dataset as *Serengeti*.

#### Snapshot Safari 2024 Expansion

This dataset aggregates camera trap observations from 15 sites across South and East Africa (Pardo et al. 2021). It includes 4.03 million images from 2.39 million sequences captured at 1,824 camera trap locations. Like *Serengeti*, annotations are at the sequence level, and the dataset includes 151 categories, including all classes from *Serengeti*. 3.72% of the non-empty sequences are annotated with multiple labels. We refer to this dataset as *Safari2024*.

For both datasets, we use MegaDetector (Beery et al. 2019a) results provided on LILA (MDv1000-redwood for *Serengeti*, MDv5a for *Safari2024* ), with the repeat detection elimination (RDE) process that has removed many false positive detections. For each animal detection with a confidence score above 0.5, we extract the smallest square crop that includes the bounding box of the animal. All cropped images originating from the same full-image sequence inherit its label and are collectively referred to as a set of crops. Sequences labelled as empty (75.86% of *Serengeti*, 65.91% of *Safari2024* ) are excluded from this process.

We train our models using the *Serengeti* dataset, focusing on the same 46 species as in Norouzzadeh et al. (2018). Images from 179 camera trap locations are used for training, and those from 46 other locations are kept for testing. For out-of-distribution testing, we use the *Safari2024* dataset with all images corresponding to the same 46 species.

### 2.2 Multi-species sequences

Because labels were assigned at the sequence level, for sequences annotated with multiple species, it is not possible to reliably associate individual crop images with specific species. For instance, if a sequence of full images is labelled as “Zebra-Wildebeest”, we know that the two species are present in the sequence, but we cannot determine which crop corresponds to a zebra and which crop corresponds to a wildebeest. This highlights an inherent limitation of sequence-level annotation, which is often used for camera trap projects: it does not provide the per-crop labels required for standard supervised learning. Furthermore, as we discuss in Section 2.3.1, this uncertainty can extend to sequences labelled as mono-specific, where unlabelled background animals may be detected but not annotated. Consequently, all our sets of crops derived from multi-species sequences can be considered weakly labelled for our task: they cannot be used for supervised training or quantitative evaluation at the crop level. Therefore, to be able to train and evaluate our model with multi-species sets of crops, we had to create synthetic ones using only the mono-specific sets. The creation process is described below.

#### Synthetic sets

We generate synthetic sets by concatenating two or three mono-specific sets of crops captured within the same hour of the day (but not necessarily on the same date). More precisely, to create synthetic sets, we first group together all mono-specific sets captured during the same hour of the day. Then, for each group, we randomly sample, without replacement, pairs or triplets of sets and concatenate them to form synthetic sets. During training, we repeat the process at each training epoch to create new combinations and increase the diversity seen by the model. Synthetic sets may not all be multi-specific, as the procedure may concatenate sets with the same species in. This is actually good for training purposes as it allows the model to learn how to use contextual information also in the monospecific case.

For evaluation, we adopt a more realistic variant of this approach: groups are formed with sets captured during the same hour, within the same month and year, and at the same camera trap location. This results in synthetic sets that share the same original background, as well as similar vegetation and lighting conditions. Evaluation on these more realistic synthetic sets therefore provides a better approximation of the real-world performance. However, these more realistic synthetic sets were not used for training as the additional constraints reduce the number of possible combinations, and thus the diversity that is important for encouraging generalisation. The statistics of the different sets are available in the Supporting Information Table 1.

**Table 1:** Accuracy (%) of the two methods for the two test sets based on the real crop sets (always mono-specific). Bold value indicate the best method. Accuracy is computed at the crop-level.

| Model | Serengeti | Safari2024 |
| --- | --- | --- |
|  | Mono-sp | Mono-sp |
| Independent Classifier | 90.48 | 92.64 |
| MCA Classifier | **95.25** | **95.02** |

Real multi-species crop sets, although weakly annotated at the crop level, can still be used for evaluation in two complementary ways. First, we can qualitatively assess them through visual inspection associated with the model’s predictions, providing insight into the model’s performance in non-synthetic multi-specific scenarios. Second, although individual crop labels are unknown, the set of species known to be present is available. As sets do not contain duplicates, each species counts only once, even if it appears multiple times in the crops. For example, a set of ten crops can be labelled {Zebra, Wildebeest}. Thus, we can evaluate them in the context of a multi-label classification task at the set level: for each set of crops, we want to be able to predict as accurately as possible the set of species known to be present.

### 2.3 Model

#### 2.3.1 Independent Classifier baseline approach

To evaluate the benefit of using our proposed new model architecture, we first have to define the state-of-the-art baseline approach against which we will compare. We naturally select to use the species classification approach commonly used in many existing platforms such as PyTorch-Wildlife, DeepFaune or AddaxAI (Hernandez et al. 2024; Rigoudy et al. 2023; Lunteren 2023) as baseline, naming it the “Independent Classifier” baseline approach. In this approach, animals are first detected in the camera trap image or sequence using a dedicated object detector (often MegaDetector v5 or v6). Each detected animal is then individually cropped and classified using a species recognition model (Figure 1). This method has proved to be very effective (Norman et al. 2023; Gadot et al. 2024). However, since each crop is processed independently, this approach cannot directly leverage the contextual information available within the set.

**Figure 1.**
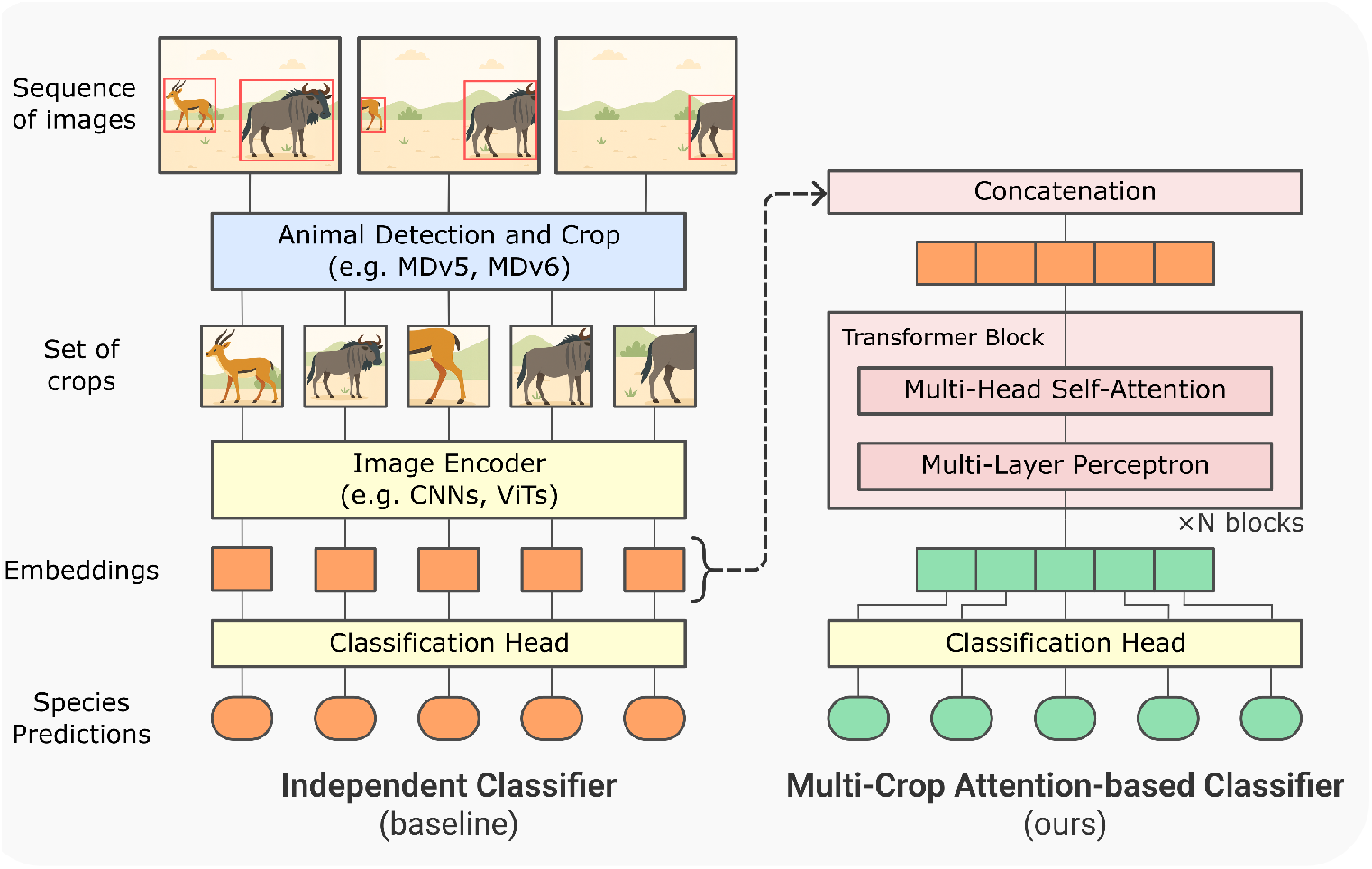
Overview of the proposed methods. A sequence of full images is first processed by an object detector, which predicts animal bounding boxes that are used to generate cropped images, each containing a single animal. A pretrained species classification model is divided into two parts: the image encoder and the classification head. Each crop is passed through the image encoder to extract feature embeddings. In the Independent Classifier baseline (left), embeddings are individually classified using the classification head. In the Multi-Crop Attention-based (MCA) Classifier method (right), embeddings are concatenated and passed through multiple transformer blocks. The resulting context-enriched embeddings are then classified using the same classification head.

To train the baseline classifier, we fine-tune the state-of-the-art DINOv2-Large model (Oquab et al. 2023), which uses the Vision Transformer (ViT) architecture (Dosovitskiy et al. 2021). The model is trained using the timm library (Wightman et al. 2023). Training hyperparameters are detailed in Supporting Information Table 3.

Since training relies on crops extracted from sequences annotated as mono-specific, all crops within a set are expected to be of the same species. However, we observe a significant number of mislabelled crops. In some cases, full images labelled with a single species also contain additional species in the background. For example, an image labelled “Wildebeest” might feature a wildebeest prominently in the foreground but also include zebras far in the background. These background animals can be detected by MegaDetector, resulting in mislabelled crops. To address this issue, we use the trained classifier to identify crops that are likely mislabelled. A crop is flagged as a likely mismatch if either (i) a class different from the ground-truth is predicted with a score greater than 95%, or (ii) the ground-truth class is predicted with a score lower than 5%. Crops identified as likely mismatches, representing 4.04% of the training data, are removed to create a “cleaned” training set. Examples of such mismatches are provided in the Supporting Information Figure 1. We then train a second classifier on this cleaned dataset, using the same architecture and hyperparameters as the original model. Importantly, no crops are removed from the test sets. This second classifier trained on the cleaned dataset is used for all subsequent experiments.

#### 2.3.2 Multi-Crop Attention-based (MCA) Classifier

We propose a simple, yet effective approach to process the set of crops jointly while reusing as many components as possible from the Independent Classifier baseline (Figure 1).

Most classification models, such as CNNs and Vision Transformers (ViTs), can be divided into two parts: an image encoder that processes the input crop image *I* to produce an embedding *e*, and a classification head (typically a single linear layer with a softmax activation) that maps the embedding to the final classification scores *s* :

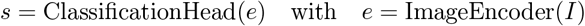

In the Multi-Crop Classifier, the architecture remains identical to the one of the Independent Classifier up to the embedding stage. However, instead of passing embeddings directly to the classification head, they are first enriched with the context provided by the set of crops. Specifically, embeddings from all crops in the set are concatenated and processed through multiple transformer blocks to produce new context-aware embeddings. These updated embeddings are then passed independently through the classification head to generate final predictions for each crop. It is worth noting that no positional embeddings are added, making the model invariant to the order of the crops within the set.

The self-attention mechanism used within the transformer blocks allows each crop embedding to pay attention to all others in the set. Here, it enables the model to capture relationships with other detections in the image or in other images of the sequence. This means that the contextual information used by the model can be the individual itself in other images or other individuals in the image or others from the same sequence. Specifically, given a set of embeddings *E* = (*e*_1_, *e*_2_, …, *e*_*N*_ ) ∈ ℝ^*N×d*^, where *N* is the number of crops in the set and *d* the embedding dimension, the transformer computes queries, keys, and values, each of dimension *d*_*k*_ with:

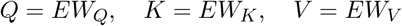

where 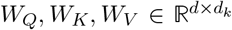 are learnable weight matrices. Using the dot product between the query and the keys and the softmax function applied on the rows, we obtain the attention matrix *A* ∈ [0, 1]^*N×N*^ :

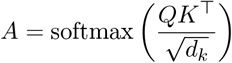

where the softmax is applied row-wise so that each row of *A* sums to 1. Each element *A*_*ij*_ can be seen as how much the crop *i* pays attention to the crop *j*. New embeddings 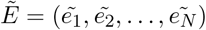 are then obtained by adding the original embeddings with a weighted sum of the attention weights and the values:

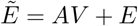

These are then passed through a Multi-Layer Perceptron (Goodfellow et al. 2016), here two linear layers, and another residual connection (He et al. 2016):

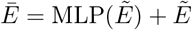

After multiple transformer blocks, these new context-enriched *Ē* crops are then processed by the classification head.

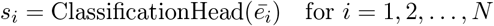

The explanation above describes the mechanism with a single self-attention head. In practice, multiple attention heads are used, but the process remains essentially the same. Additionally, each step includes a layer normalization, that was omitted in the description above for clarity. Interested readers are referred to the original paper by Vaswani et al. (2017) for more details.

During training, both the image encoder and classification of the pretrained classifier are kept frozen. For a fair comparison, we use the same classifier trained in the previous section, although any existing pretrained classifier could be used. Crop embeddings are precomputed once and saved in a memory bank, only the transformer blocks are trained, directly from the embeddings.

In our experiments, the Multi-Crop Classifier is trained using four transformer blocks, a batch size of 512, the AdamW optimizer (Loshchilov et al. 2019), and a learning rate of 2 *×* 10^−6^. During training, all sets within a batch must have the same length, and here we use a fixed set length of 12 crops. Longer sets are randomly downsampled to fit this constraint and shorter sets are discarded. This limitation applies only during training, as at inference time this architecture can handle sets of arbitrary length.

Since the Multi-Crop Classifier is trained using embeddings derived from the crop used to train the Independent Classifier, these embeddings are more “overfitted” than if they originated from unseen crops and can create a distribution shift. To improve robustness, we apply embedding mixup (Zhang et al. 2018; Verma et al. 2019), a regularization technique in which an embedding *e*_*i*_ is mixed with a randomly selected embedding *e*_*r*_ from the memory bank, thereby reducing the amount of information it carries. The mixed embedding is computed as:

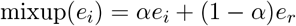

We use a mixup factor *α* = 0.5 for one third of the set, and a lower *α* = 0.8 for the remaining two thirds. These values were chosen empirically to balance regularization strength, while ensuring that the embeddings still carry meaningful information.

### 2.4 Metrics and experiments

We compare the two methods (Independent and MCA classifiers) using accuracy as the primary evaluation metric. Accuracy is always computed at the crop level, regardless of whether crops are processed individually or within sets. It corresponds to the proportion of crops correctly classified:

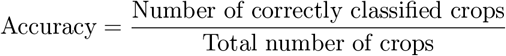

We evaluate both methods on original and synthetic sets. For the original sets, we retain only those that have least two crops (87.00% for *Serengeti* and 67.35% for *Safari2024* ).

For the real, but weakly annotated, multi-specific sets of crops, it is not possible to measure accuracy since labels are not available at the crop level. Hence, we use the Jaccard index, a standard metric for multi-label classification (Sorower 2010). Given, a ground-truth set of labels *G* and a predicted set of labels *P* the Jaccard index J is the size of their intersection divided by the size of their union:

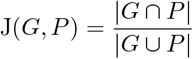

To construct the predicted label set *P*, we classify each crop and retain only the labels whose confidence score is above a given threshold. We then average the Jaccard index across all sets of crops and report the performance of the model at different threshold values. For each threshold, we also report how many crops were excluded due to their prediction scores falling below the threshold.

We use the Expected Calibration Error (ECE) to measure the calibration of the scores produced with the two models (Guo et al. 2017; Dussert et al. 2024). A perfect calibration would be achieved when the ECE is equal to 0, as in this case the scores can be interpreted as the probability of the prediction being true. We calibrate the models on the *Serengeti* test set using temperature scaling and report the ECE on *Safari2024* using the same temperature.

To provide a reference point, we also compare our baseline against SpeciesNet (Gadot et al. 2024), a publicly available state-of-the-art species classification model for cameratrap images. For this comparison, we applied SpeciesNet version 4.0.1a directly to the already cropped images. Since SpeciesNet has a larger set of classes than our model, we mapped its classes to the one used in this study, using a procedure detailed in Supporting Information Section 4. We report SpeciesNet accuracy only for *Safari2024* since the test set of *Serengeti* used in this study is included in SpeciesNet training set.

We also perform an ablation study to assess the impact of our training design choices by comparing the main configuration to alternative setups in which only a single parameter is modified. Finally, we provide a qualitative assessment of the model using real multi-species crop sets. In particular, we visualize the attention matrix from the final transformer block, as it offers valuable insights on the model’s behaviour.

## 3 Results

Our new Multi-Crop Attention-based (MCA) Classifier reaches an accuracy above 95% on the test datasets (real, mono-specific sets), outperforming the Independent Classifier by 4.77% on *Serengeti* and 2.38% on *Safari2024* (Table 1). For comparison, SpeciesNet has an accuracy of 91.74% on *Safari2024*, which is 0.90% lower than the Independent Classifier.

The MCA Classifier also performs better on the synthetic, mono- or multi-specific sets (Table 2). While the gap is lower than in mono-specific sets, there is still a significant improvement in multi-specific sets with an improvement of 3.34% on *Serengeti* and 2.23% on *Safari2024* over the Independent Classifier.

**Table 2:** Accuracy (%) of the two methods for the two test sets on the synthetic sets. Bold value indicate the best method. Accuracy is computed at the crop-level.

| Model | Serengeti |  | Safari2024 |  |
| --- | --- | --- | --- | --- |
|  | Mono-sp | Multi-sp | Mono-sp | Multi-sp |
| Independent Classifier | 92.18 | 85.48 | 93.09 | 91.20 |
| MCA Classifier | **97.53** | **88.82** | **95.72** | **93.57** |

As real multi-species sets are only weakly labelled (see Methods), we first conduct qualitative visual analyses at the crop level. One illustrative example is shown in Figure 2 and additional cases are shown in Supporting Information Figures 1 and 2. We then evaluate both models at the set level using the Jaccard index (Figure 3). For the Independent Classifier, the Jaccard index starts at 0.796 with a threshold of 0.2, peaks at 0.860 with a threshold of 0.85, discarding 20.1% of crops, indicating it initially predicts too many species. It then declines as the threshold becomes too high, and the model begins predicting too few species. In contrast, the MCA Classifier shows minimal variation, achieving 0.863 at threshold 0.2 and a maximum of 0.864 at threshold 0.6, discarding only 2.87% of crops.

**Figure 2.**
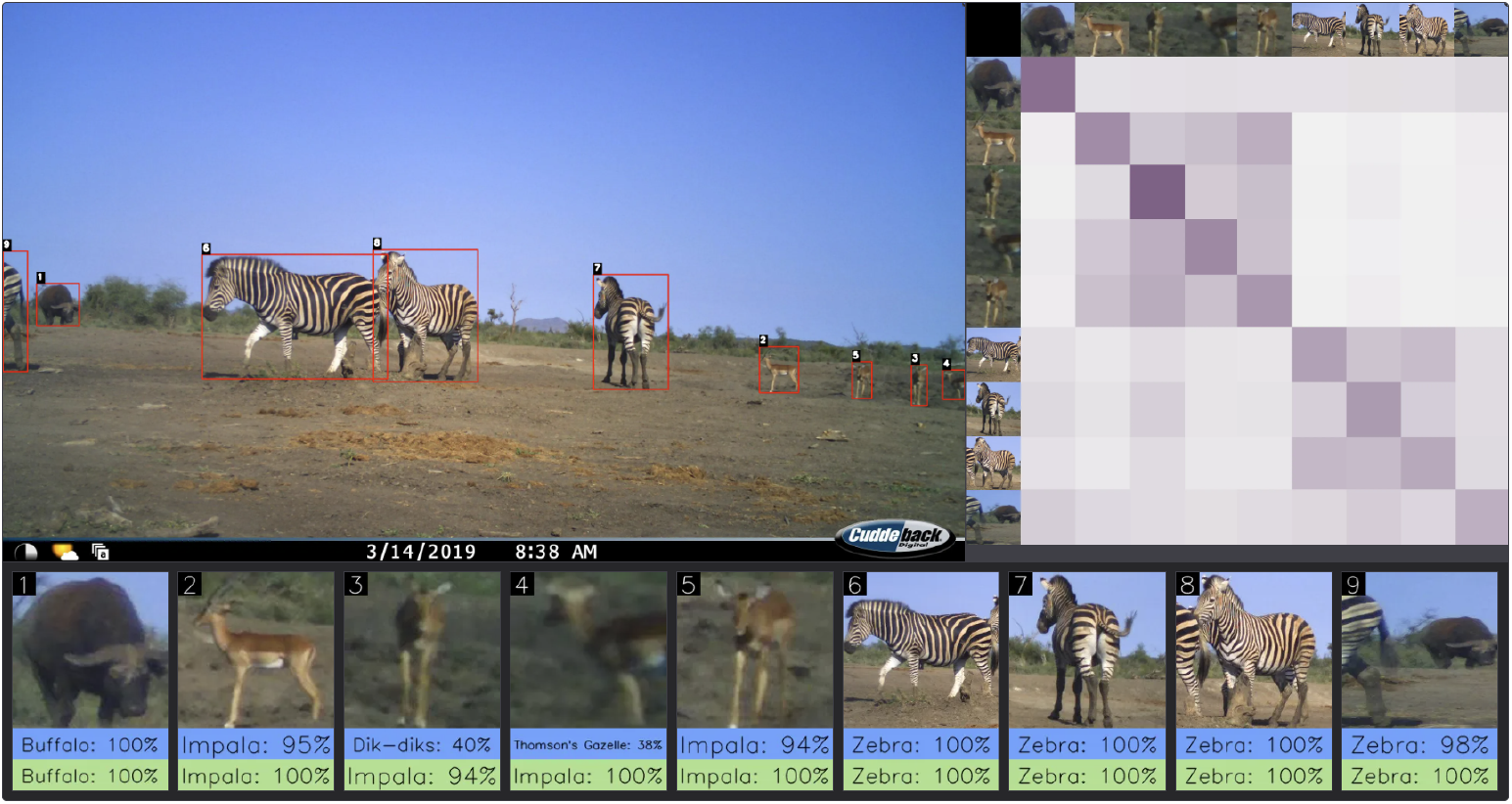
Qualitative analysis of the MCA approach on a multi-species image labelled {Impala, Buffalo, Zebra} from the *Safari2024* dataset. The top-left panel shows the original image with animal detections by MegaDetector. The bottom row displays the cropped detections with their predicted labels: blue for the Independent Classifier, and green for the Multi-Crop Attention-based Classifier. The top-right panel presents the attention matrix, where each row corresponds to one crop embedding and displays how much attention it attends to other crops in the sets (columns). The attention weights in each row always sum to 1. In the colour scale, white represents an attention weight of 0, and dark purple indicates an attention weight of 1. Here, the 3^rd^ and 4^th^ crops are misclassified by the independent classifier, but correctly identified as impalas by the self-attention model. The attention matrix indicates that they strongly attend to other impalas crops, highlighting the benefit of self-attention in this context.

**Figure 3.**
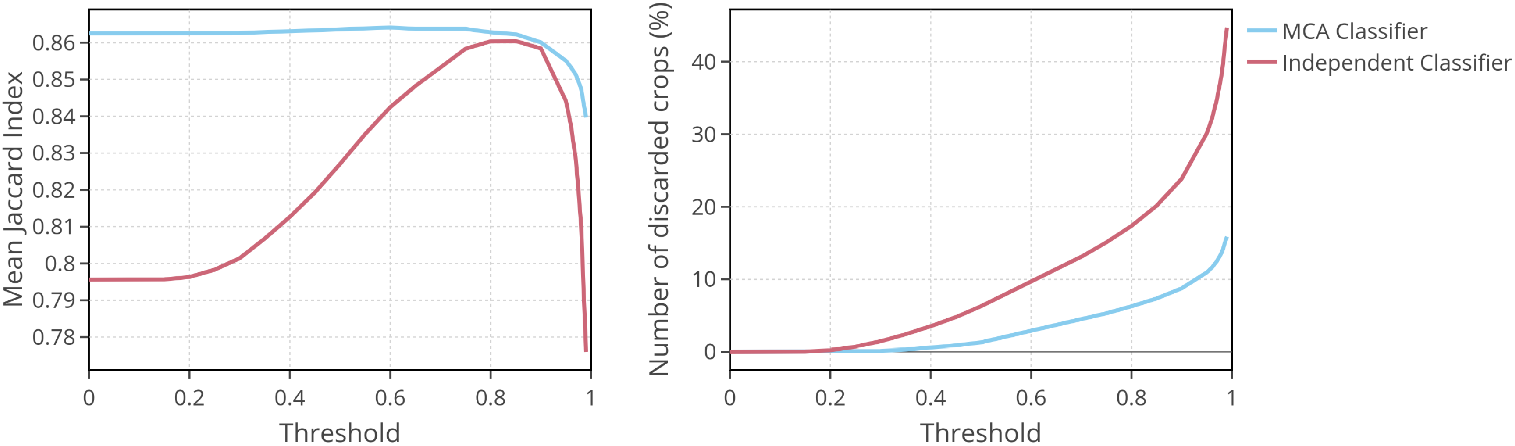
Similarity between species composition in crop sets: (left) average Jaccard index across all crop sets, using real multi-specific test sets of *Serengeti* and *Safari2024*, at different threshold values. (right) Percentage of crops discarded due to their predicted scores being below the threshold.

Table 3 presents the results of an ablation study, comparing the configuration used in the main results (shown in the last row of each table) with several alternative configurations. Training on real, but mono-specific only, sets of crops instead of synthetic ones significantly reduces multi-species accuracy, even performing worse than the Independent Classifier. Increasing the number of transformer blocks from one to two yields a small improvement, with a further minimal gain observed when using four blocks. Additionally, applying embedding mixup during training also provides an overall benefit.

**Table 3:** Ablation study on different hyperparameters of our approach. We report the accuracy for mono-specific (Mono-sp) and multi-specific (Multi-sp) synthetic sets of the *Safari2024* dataset. In each table, only one parameter varies relatively to the main configuration, used in the rest of the results, that is: training on synthetic sets, using four transformer blocks, and applying embedding mixup.

| Sets of crops type | Mono-sp | Multi-sp |
| --- | --- | --- |
| Real (mono-sp only) | **96.24** | 89.52 |
| Synthetic | 95.72 | **93.57** |

(a) Type of the crops sets used for training
| Mixup | Mono-sp | Multi-sp |
| --- | --- | --- |
| No | 95.25 | 93.29 |
| Yes | **95.72** | **93.57** |

(b) Embedding mixup during training
| #Blocks | Mono-sp | Multi-sp |
| --- | --- | --- |
| 1 | 95.25 | 93.30 |
| 2 | 95.65 | 93.49 |
| 4 | **95.72** | **93.57** |

(c) Number of transformer blocks

Assessing the calibration of the scores produced by the two approach using the Expected Calibration Error (ECE) reveals that the Independent Classifier has an ECE of 1.84% after scaling with the optimal temperature of 1.15. In contrast, the MCA Classifier reaches a lower ECE of 1.13%, but required a substantially higher temperature of 2.92. This indicates that while the MCA Classifier provides more reliable scores after calibration, it is initially more overconfident than the Independent Classifier.

While the MCA Classifier improves classification accuracy, it is important to quantify its computational cost. The additional transformer layers train efficiently, requiring 65 seconds per epoch and totalling 4 hours and 10 minutes on a single NVIDIA V100 GPU. At inference time, the MCA Classifier approach introduces negligible overhead, with an average slowdown of only 0.64% compared to the Independent Classifier. In these models, most of the computations are done by the Image Encoder, which is shared by both approaches.

## 4 Discussion

This work addresses a long-standing question in automated camera trap image analysis: how can we replicate the human ability to integrate temporal contexts as well as information from other animals when identifying species in camera trap images? Our proposed solution is both simple to implement and effective. We show that the self-attention mechanism (Vaswani et al. 2017), central to the success of large language models (LLMs), can be effectively used in a camera trap species classification pipeline without requiring complex implementation or labour-intensive annotations. By applying transformer blocks directly on the classifier’s embeddings, we are able to generate new representations that leverage the information contained across the set. This approach not only achieves a substantial improvement over the baseline but also has learned to handle multi-species sets, where contextual information must be interpreted carefully to preserve species-specific information.

Figure 2 and Figure 3 illustrate well a common issue that arises when a label is predicted independently for each crop: including all predictions increases the risk of false positives: predicting species that are not actually present, while filtering out low-confidence predictions can lead to underestimating the number of individuals of each species. Our approach provides, at virtually no cost, greater robustness in the face of this trade-off, resulting in more accurate estimations for downstream ecological analyses (Whytock et al. 2023). In addition, it can improve the user’s trust in the predictions of the model by reducing these false positives (Gadot et al. 2024). Moreover, attention maps produced by the self-attention layers can offer some interpretability (Dosovitskiy et al. 2021), helping to visualize which crops the model relies on for its predictions.

While the MCA Classifier achieves higher accuracy and a lower ECE after calibration, the requirement for a high temperature scaling factor (*T* = 2.92 vs *T* = 1.15 for the baseline) indicates that the model’s raw predictions are significantly overconfident. We attribute this to the nature of the self-attention mechanism pooling information across multiples crops which can amplify consistent patterns: when many crops support the same interpretation, the model becomes increasingly confident in its prediction. This behaviour highlights a critical consideration for deployment: raw confidence scores from the model should not be interpreted as direct probabilities. Although temperature scaling effectively corrects this in post-processing, future work could investigate strategies for addressing this during training, such as label smoothing or mixup (Szegedy et al. 2015; Zhang et al. 2018).

In this work, we chose to process all the crops from a given sequence simultaneously. However, the architecture could technically use more context, for example, by adding the crops from previous or following sequences in a given time interval. Yet, by expanding the context, there is a risk of introducing unintended biases into the model’s predictions, especially when the added context is not directly relevant to the target crops (Shi et al. 2023; Liu et al. 2023). This raises an important question: What is the appropriate amount and type of context to provide to the model? A reasonable answer may be to match the context available to human annotators. In our case, both datasets used in this study were labelled on Zooniverse, where participants had access to the entire sequence produced by the camera trap during annotation.

Although the ability to train the classifier and the transformer blocks separately can be seen as a strength of the approach, enabling modularity and efficiency, it may also limit the overall expressiveness of the architecture. Since the transformer blocks are trained on fixed embeddings, the transformer’s capacity to refine the embeddings is constrained by what the classifier has learnt. Joint training of the classifier and transformer could potentially achieve better performance on crop sets by allowing end-to-end optimization. However, standard joint training would substantially increase GPU memory requirements, particularly with large batch sizes, and may limit the ability of practitioners with limited computational resources to train the model, thereby reducing its accessibility for broader adoption and experimentation. Nevertheless, future work could explore strategies to enable partial end-to-end optimization. For instance, freezing the earlier layers of the image encoder or using Low-Rank Adaptation (LoRA, Hu et al. 2021), could allow the model to refine feature extraction specifically for the multi-crop context, without requiring full fine-tuning and its associated prohibitive memory costs.

In this study, we trained both the species classification model and the transformer blocks that operate on its crop embeddings. However, in practice, pretrained classifiers are already available, such as in SpeciesNet or DeepFaune (Gadot et al. 2024; Rigoudy et al. 2023), and only the transformer module would need to be trained, provided that training data (either images or precomputed embeddings) are available. Nonetheless, our approach has two main limitations. First, the training data must include sequence identifiers, i.e. a unique ID indicating which crops belong to the same sequence. Second, the training data must cover all species classes present in the classifier. Requiring sequence identifiers can reduce the amount of training data that can be used, compared to the traditional Independent Classifier, as such identifiers are not always provided. Since image metadata typically does not include sequence information, additional effort is needed to obtain it, either through manual annotation or by inferring it from timestamps and location data (if available)(Bubnicki et al. 2024).

This new approach helps to address some of the blind spots in the species classification pipeline by improving performance under challenging conditions such as poor crop quality or occlusion (e.g., Figure 2). However, these improvements alone are not sufficient to reach perfect accuracy, as other limitations persist elsewhere in the pipeline. For instance, even the most accurate animal detectors (e.g., MDv5a, MDv6-YOLOv10-Extra, Beery et al. 2019a; Hernandez et al. 2024) occasionally fail to detect animals, particularly in cases involving small or well-camouflaged species (Brookes et al. 2024; Mulero-Pázmány et al. 2025). In contrast, human annotators can often identify such animals by relying on their movement across images in a sequence. Thus, the development of new generalist animal detection models that, like humans, can leverage the motion to detect animals, could bring us closer to fully automating species identification in camera trap data. This would be another way to handle temporal context, this time for detection purposes.

Finally, while we demonstrated the benefits of this approach for species classification, it could also be adapted to other computer vision tasks that can benefit from temporal context and information from surrounding animals, such as behaviour classification or individual counting. Many models developed for video data rely on short and regular frame intervals (Feichtenhofer et al. 2017; Karaev et al. 2023) and tend to perform poorly on camera trap sequences, where time gaps between images can be of several tens of second. In contrast, with our approach, applying self-attention to crop embeddings offers a simple and flexible solution, as it can learn to capture the relationship between crops without temporal continuity.

## 5 Conclusion

We introduced a lightweight self-attention module that improves species classification in camera trap images by using contextual information from other animal detections within the same event. Our method integrates seamlessly into existing two-step pipelines, handles multi-species settings, and with minimal computational cost. By reducing species misclassifications, it enables more reliable estimates of biodiversity metrics, such as species richness, occupancy, and interspecies interactions, which are essential for wildlife monitoring and conservation

## Acknowledgements

This work was granted access to the HPC resources of IDRIS under the allocation 2022-AD010113729 made by GENCI.

## Author contributions

Gaspard Dussert, Simon Chamaillé-Jammes, Stéphane Dray and Vincent Miele conceived the ideas and designed the methodology. Gaspard Dussert coded and performed the analysis. Gaspard Dussert wrote the first version of the manuscript, Simon Chamaillé-Jammes, Stéphane Dray and Vincent Miele contributed critically to the drafts and gave final approval for publication.

## Data availability statements

The code will be made available on GitHub : https://github.com/gdussert/MCA_Classifier

The derived data used in the analyses will be made available on Zenodo : https://zenodo.org/records/15736090

## Supporting information

## 1 Datasets statistics

**Table 1:** Crop sets statistics of the different datasets. * indicates that the crops sets are weakly labelled and cannot be used for supervised training or evaluation at the crop level (per-crop accuracy).

| Dataset | Type | Mono-sp |  | Multi-sp |  |
| --- | --- | --- | --- | --- | --- |
|  |  | #Sequences | #Crops | #Sequences | #Crops |
| *Serengeti* (train) | Real | 440k | 4.13M | 21.0k* | 441k* |
| *Serengeti* (train) | Synthetic | ~27k | ~319k | ~153k | ~1,83M |
| *Serengeti* (test) | Real | 163k | 1,49M | 6.92k* | 137k* |
| *Serengeti* (test) | Synthetic | ~131k | ~2,25M | ~63.6k | ~739k |
| *Safari2024* (test) | Real | 301k | 1,24M | 18.1k* | 219k* |
| *Safari2024* (test) | Synthetic | ~244k | ~1.83M | ~107k | ~657k |

**Table 2:**
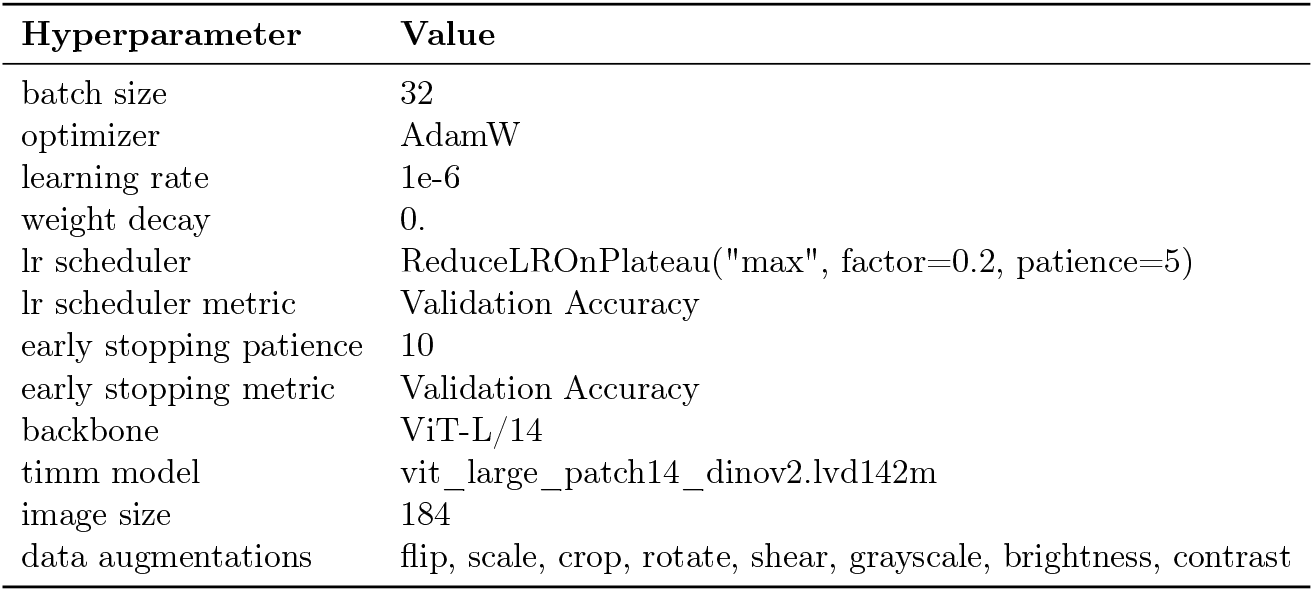
Crop Classifier training hyperparameters.

## 2 Training hyperparameters

**Table 3:** Multi-Crop Attention-based Classifier training hyperparameters.

| Hyperparameter | Value |
| --- | --- |
| batch size | 512 |
| optimizer | AdamW |
| learning rate | 2e-6 |
| weight decay | 0. |
| lr scheduler | ReduceLROnPlateau("min", factor=0.2, patience=15) |
| lr scheduler metric | Validation Loss |
| early stopping | lr < 1e-8 |
| transformer blocks | 4 |
| attention head | 8 |
| dim | 1024 |
| mlp dim | 4096 |
| normalization | Layer Norm |
| activation | GELU |
| data augmentations | None |

## 3 Additional figures

**Figure 1.**
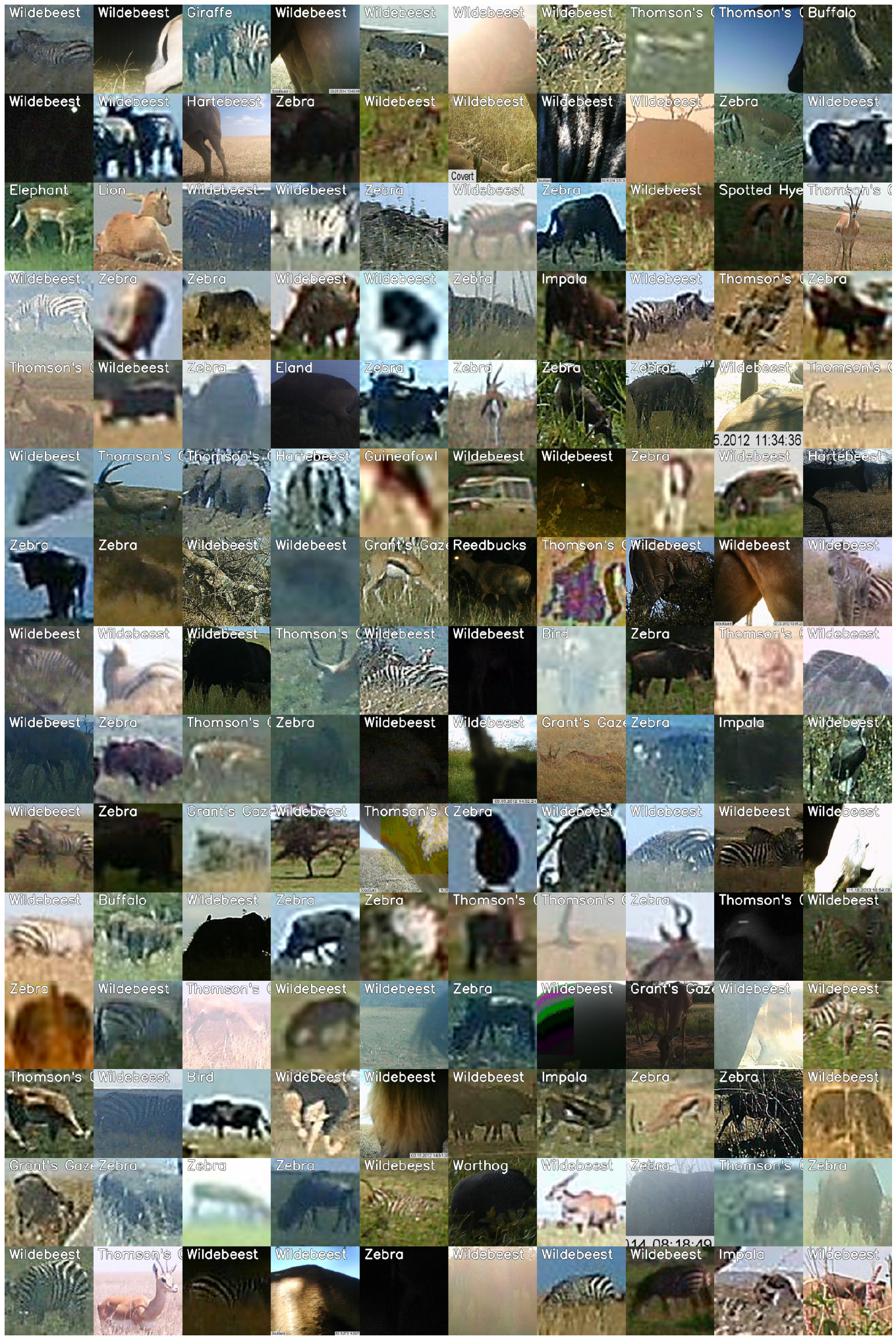
Examples of likely mismatches (random sample) that were excluded from the second training of the classifier.

**Figure 2.**
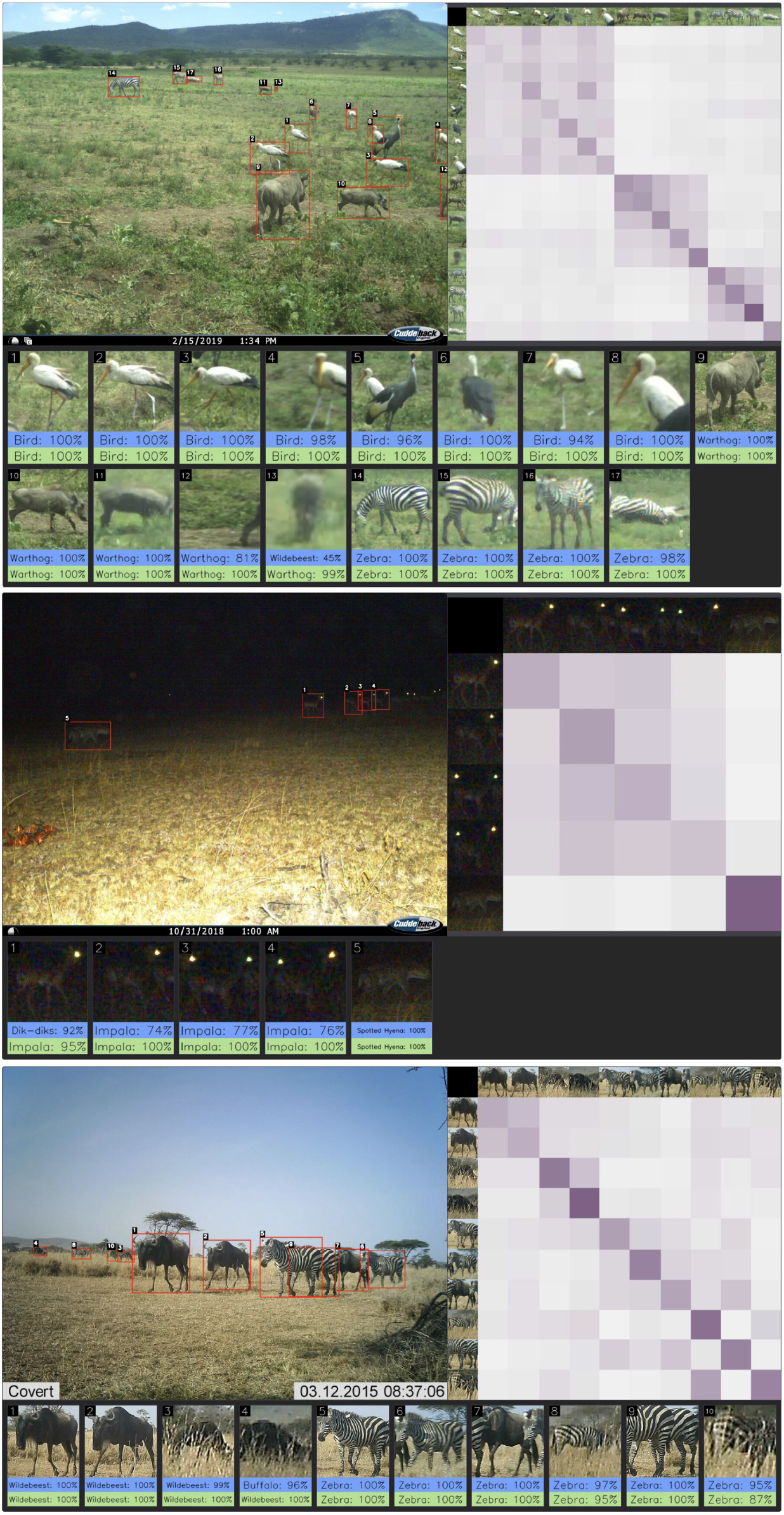
Additional predictions of the two approaches, on test multi-species images (single picture): blue for the Independent Classifier, and green for the MCA Classifier. From top to bottom, the images are labelled “Bird & Warthog & Zebra”, “Impala & Spotted Hyena” and “Wildebeest & Zebra”

**Figure 3.**
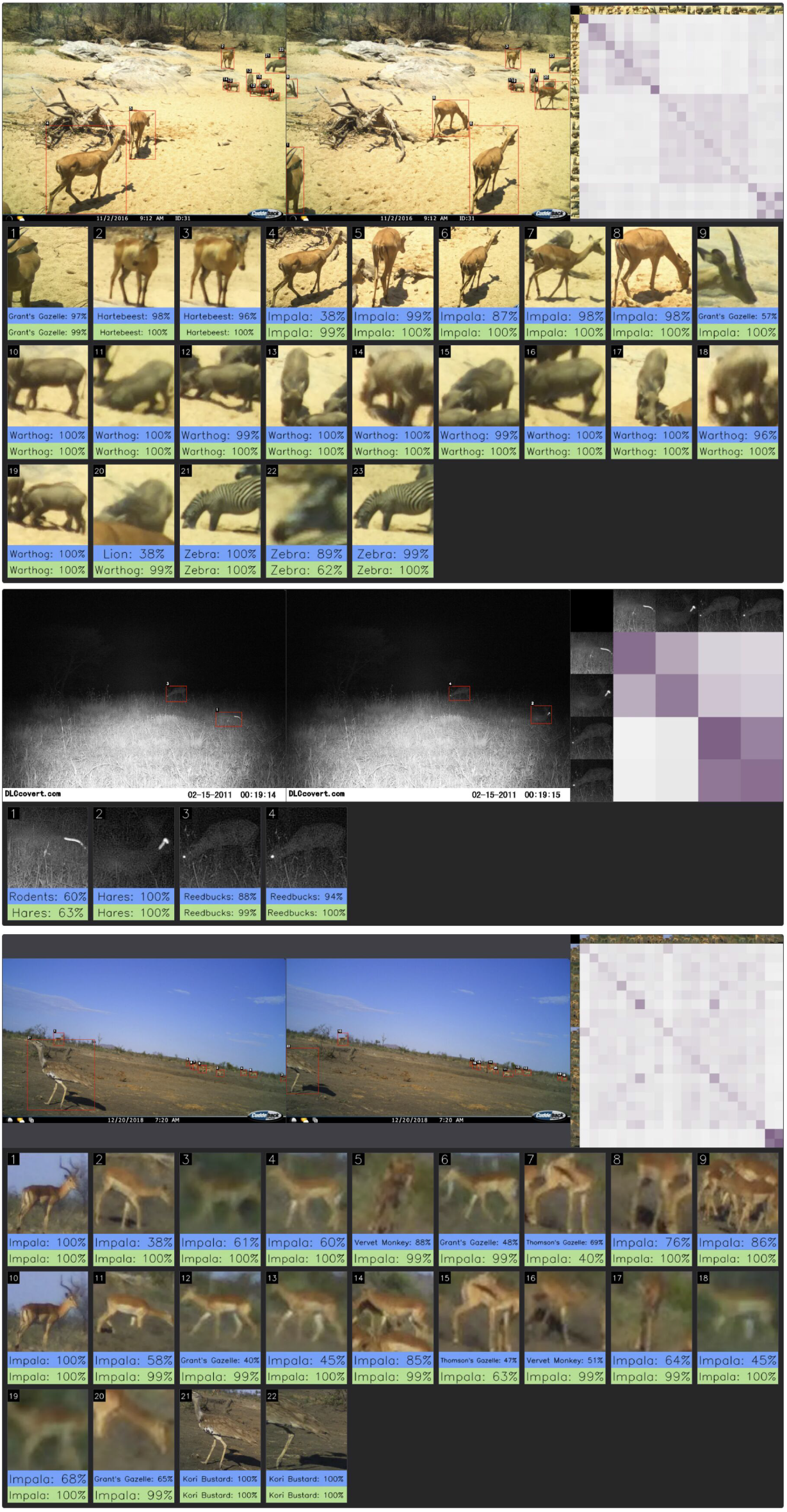
Additional predictions of the two approaches, on test multi-species images (sequence of two pictures): blue for the Independent Classifier, and green for the MCA Classifier. From top to bottom, the images are labelled “Impala & Hartebeest & Warthog & Zebra”, “Dik-Dik & Hare” and “Impala & Kori Bustard”

## 4 Comparison with SpeciesNet

To provide a comparison with a publicly available model, we used the state-of-the-art classification model, SpeciesNet (Gadot et al. 2024). We used the speciesnet python library (version 5.0.2) with the SpeciesNet 4.0.1a model in “classifier-only” mode, as the image crops had already been generated.

Since SpeciesNet has a larger set of classes than our model, we had to map its predicted classes to the list of species used in our study for comparison. For example, any *Lagomorpha* species predicted by SpeciesNet was converted to the “hare” class (since it is the only *Lagomorpha* species in the classes of the study).

To handle SpeciesNet classes that could not be directly mapped (e.g., “blank” or “*Bos taurus*”, which would result in an incorrect prediction), we employed the following strategy: we computed the scores for all SpeciesNet classes, sorted them from highest to lowest, and selected the class with the highest score that could be successfully mapped to one of our model’s classes.

The SpeciesNet predictions are available in the data release, and the scripts for class mapping and accuracy computation are available in the code release.

## 5 Sequence aggregation of predictions for mono-specific sequences

When sequences are known to be mono-specific, post-processing methods can be employed to aggregate per-crop predictions into a unified sequence prediction. In this section, we investigate the impact of such aggregation on the real (non-synthetic) sequences of *Serengeti* and *Safari2024*. To this end, we use the Average Logit method, which has been shown by Dussert et al. (2024) to provide both accurate and calibrated sequence-level predictions. This method averages the predicted logits across the different crops before normalizing them with the softmax function, to obtain sequence-level scores for each species. Each crop in the sequence is then classified as the species with the highest score.

As shown in Table 4, the Average Logit method improves performance for both the Independent Classifier and the MCA Classifier, and reduces the gap between the two models. While efficient, it is crucial to note that this post-processing strategy enforces a single species label per sequence. Consequently, it is not suitable for multi-specific sequences and would be difficult to implement in habitats where multi-species events are frequent, as there is no prior knowledge of whether a sequence is mono- or multi-specific.

**Table 4:**
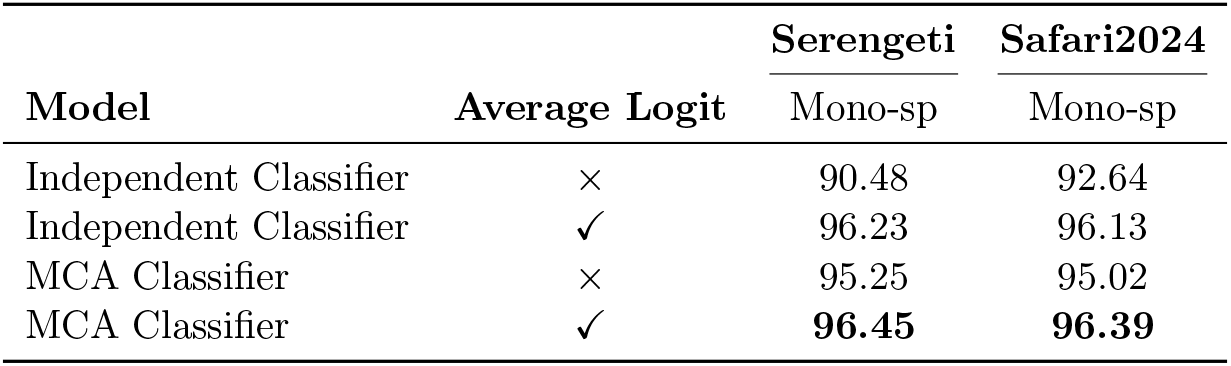
Accuracy (%) of the two methods for the two test sets (*Seregenti* and *Safari2024* ) based on the real crop sets (always mono-specific), with and without Average Logit, a post-processing aggregation method. Bold value indicate the best method. Accuracy is computed at the crop-level.

| Model | Average Logit | Serengeti | Safari2024 |
| --- | --- | --- | --- |
|  |  | Mono-sp | Mono-sp |
| Independent Classifier | × | 90.48 | 92.64 |
| Independent Classifier | ✓ | 96.23 | 96.13 |
| MCA Classifier | × | 95.25 | 95.02 |
| MCA Classifier | ✓ | **96.45** | **96.39** |

